# iHypoxia: an integrative database of the expression dynamics of proteins in response to hypoxia in animals

**DOI:** 10.1101/2021.10.16.464637

**Authors:** Ze-Xian Liu, Panqin Wang, Qingfeng Zhang, Shihua Li, Yuxin Zhang, Yutong Guo, Chongchong Jia, Tian Shao, Lin Li, Han Cheng, Zhenlong Wang

## Abstract

Decreased oxygen concentrations (hypoxia) outside of the physiological range may severely subvert cell, tissue, and organism survival. Mammals have evolved mechanisms to sense hypoxia and induce a series of hypoxic responses. In recent years, high-throughput techniques have greatly promoted global perturbation studies of protein expression during hypoxia, and these studies have contributed to the understanding of the complex regulatory networks of hypoxia. In this study, we developed an integrated resource for the expression dynamics of proteins in response to hypoxia (iHypoxia), which contains 1,629 expression events of 1,215 proteins identified by low-throughput experiments (LTEs) and 154,953 quantitative expression events of 36,194 proteins identified by high-throughput experiments (HTEs) from five mammals that exhibit a response to hypoxia. Various experimental details such as the hypoxic experimental conditions, expression patterns, and samples were carefully collected and integrated. In addition, we conducted an orthologous search and identified 581,763 proteins that may respond to hypoxia among 50 animals. An enrichment analysis of human proteins identified from LTEs showed that these proteins were enriched in certain drug targets and cancer genes. The annotation of known posttranslational modification (PTM) sites to proteins identified by LTEs revealed that these proteins underwent extensive PTMs, particularly phosphorylation, ubiquitination and acetylation. Based on the results, iHypoxia provides a convenient and user-friendly method for users to obtain hypoxia-related information of interest. We anticipate that iHypoxia, which is freely accessible at http://ihypoxia.omicsbio.info, will advance the understanding of hypoxia and serve as a valuable data resource.

## Introduction

Oxygen is an essential element for the survival of aerobic organisms, and a decreased oxygen concentration (hypoxia) may threaten the survival of cells, tissues and individuals. To cope with hypoxia, mammals have evolved sophisticated oxygen-sensing systems and adaptive mechanisms [1], including erythropoiesis [2], mitochondrial respiration [3], angiogenesis [4] and metabolic adaptations [5]. In 1995, Wang *et al*. identified hypoxia-inducible factor 1 (HIF-1), which is a heterodimer consisting of a constitutively expressed HIF-1β subunit and a HIF-1α subunit that is regulated and expressed in an oxygen-dependent manner [6]. Under normoxic conditions, HIF-1α is hydroxylated by prolyl hydroxylases and then targeted for proteasomal degradation through Hippel-Lindau (VHL)-mediated ubiquitination [7-9]. During hypoxia, accumulating HIF-1α dimerizes with HIF-1β and then binds to hypoxia response elements to trigger the hypoxic response [10]. In addition to HIF-1, many proteins and pathways are involved in hypoxia signaling [11] and thus play important roles in various physiological and pathological processes, such as the regulation of immunity [12], embryonic development [13], cancer [14], cardiovascular diseases [15], and anemia [16]. The use of hypoxia-related strategies for the treatment of disease is being investigated. For example, Chen *et al*. found that roxadustat, an oral inhibitor of HIF prolyl hydroxylase, is useful for treating patients with anemia and chronic kidney disease [17]. Therefore, the dissection of hypoxia signaling is critical, and an exploration of the expression dynamics of proteins in response to hypoxia is the first step.

Several databases related to hypoxia are currently available. In 2012, Pankaj *et al*. compiled HypoxiaDB, a database of hypoxia-regulated proteins containing 3,500 human proteins with detailed annotations that were identified from high-throughput experiments [18]. However, HypoxiaDB has not been updated since then. In 2017, Rashid *et al*. developed the HRGFish database, which contains 818 hypoxia-responsive genes collected from 38 fishes [19]. During the last decade, massive studies have contributed to dissecting the protein expression dynamics in cellular/animal models in response to hypoxia under various experimental/physiological conditions. Therefore, a comprehensive database that integrates the quantitative events of protein expression in response to hypoxia will boost the reutilization of previous investigations and provide helpful data resources for studying the molecular mechanism of hypoxia-related biology.

Herein, we present iHypoxia, an integrative database for the expression dynamics of proteins in response to hypoxia in animals. At present, the iHypoxia database has been curated and hosts 156,582 quantitative expression events of 36,408 proteins in five mammals in response to hypoxia and 581,763 orthologous proteins in 50 animals. Detailed annotations, including posttranslational modifications (PTMs), subcellular locations, protein-protein interactions and drug-target relations, are also provided. Search and browsing services were also implemented for user convenience. An enrichment analysis of human proteins identified in low-throughput experiments (LTEs) showed that proteins that respond hypoxia were significantly enriched in certain drug targets and oncogenes. By mapping known PTM sites to proteins identified in LTEs, we confirmed that these proteins can undergo extensive PTMs, particularly phosphorylation, ubiquitination and acetylation. We anticipate that the iHypoxia database will be beneficial to the research community, and this online service is freely available at http://ihypoxia.omicsbio.info.

## Results

### Data statistics of iHypoxia

The flow chart of the construction of iHypoxia is depicted in **Figure 1**. Through literature curation and public database integration, we obtained 156,582 quantitative expression events of 36,408 proteins that respond to hypoxia in five mammals, including *Homo sapiens* (human), *Mus musculus* (mouse), *Rattus norvegicus* (rat), *Sus scrofa* (pig) and *Bos taurus* (cow). Based on the expression alterations of the proteins in hypoxia compared with normoxia, proteins were divided into the upregulation and downregulation categories **(Figures 2A and 2B**). Because proteins that respond to hypoxia present different expression patterns in various tissues/cells or under various hypoxic conditions, some proteins may be upregulated and downregulated as conditions change and were annotated “up/down” (**Figures 2A and 2B**). In general, the protein expression quantifications obtained in LTEs were reliable, and those obtained in high-throughput experiments (HTEs) needed further validation; therefore, we curated the expression quantifications from LTEs and HTEs separately. In this study, for five mammals, we obtained 1,215 unique proteins whose expression levels showed changes in response to hypoxia detected in LTEs (**Figure 2A**) and 36,194 proteins identified by HTEs (**Figure 2B**). Regardless of whether the proteins were obtained from HTEs or LTEs, the proteins detected in humans, mice, and rats accounted for the vast majority of the proteins, and few proteins were found in cows and pigs. Furthermore, we observed that 82.39% (1001/1215) of the proteins identified in LTEs were also detected in HTEs (**Figure 2C**), and 12,290 proteins showed altered expression at both the protein and transcription levels (**Figure 2D**). The distributions of different experimental types for protein expression detection in LTEs and HTEs are shown in **Figures 2E** and **2F**, respectively. Western blotting (WB) and microarrays (arrays) were the most commonly used techniques for the detection of hypoxia-related protein expression. In addition, considering the data for one species should be helpful for studies in another species because cells might utilize the same or similar mechanisms across species, we further expanded our data to 618,171 proteins in 50 animals based on an orthologous search (Supplementary Table S1), and this expansion could provide helpful information for nonmodel animals that scientists might be interested in.

**Figure 1.**
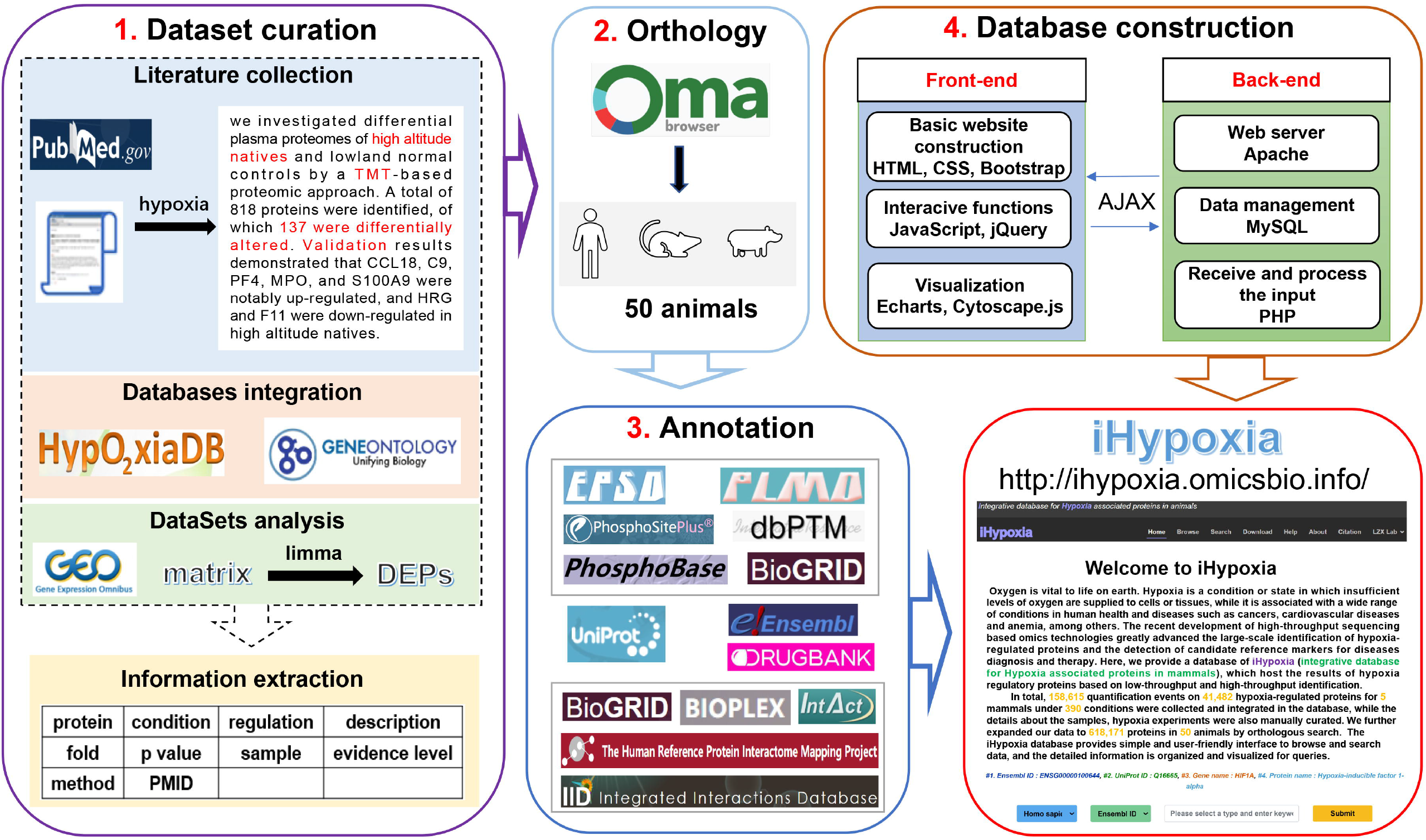
Workflow of the construction of the iHypoxia database. First, we collected and integrated expression data of proteins in response to hypoxia from the literature, HypoxiaDB, GO resources and GEO. We then performed ortholog detection using the OMA dataset. In addition to collecting basic information, we annotated proteins in four aspects: PTMs, subcellular locations, protein-protein interactions, and drug-target relations. Ultimately, the iHypoxia website was constructed.

**Figure 2.**
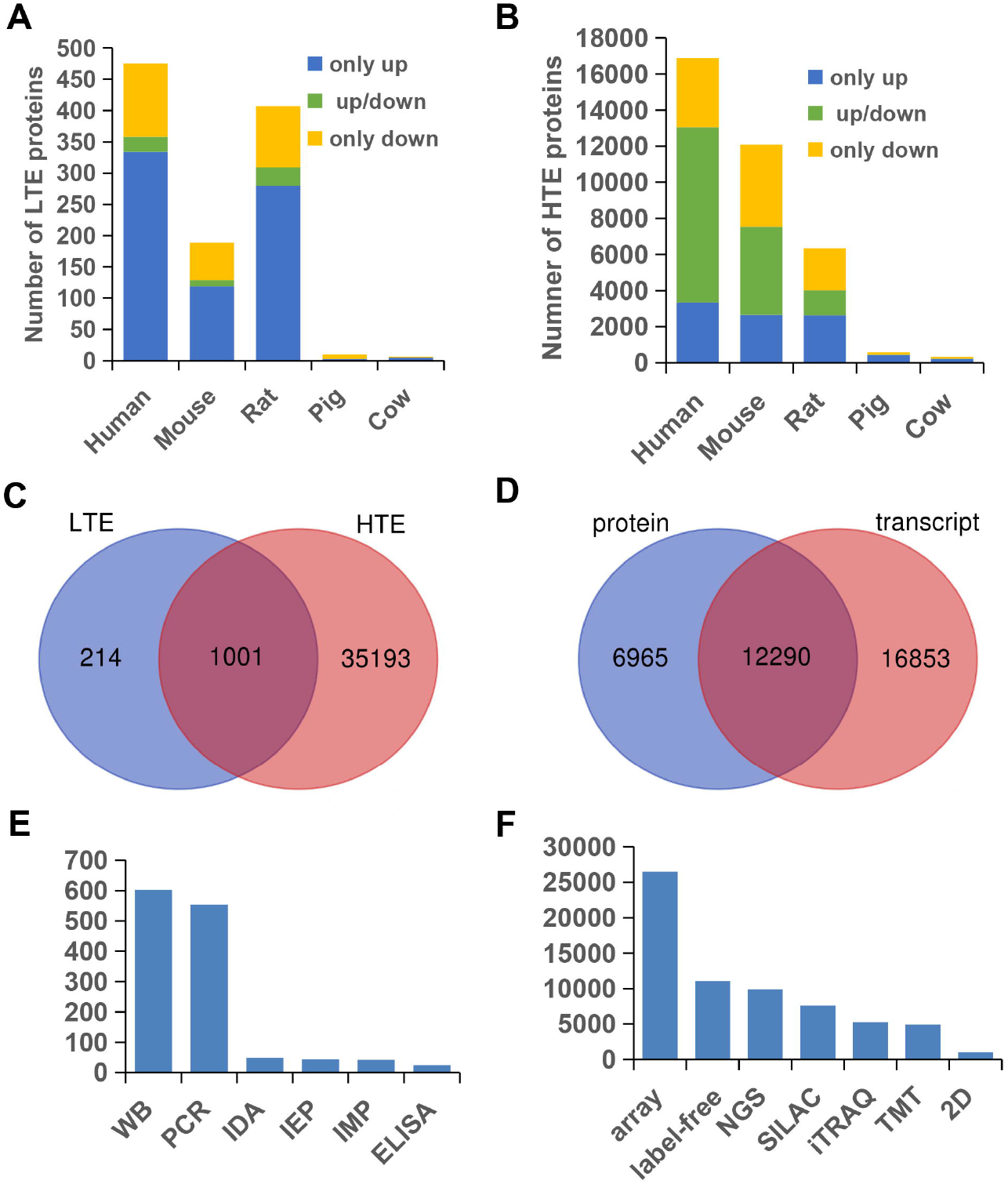
Data statistics of iHypoxia. **A**. Distribution of proteins identified from LTEs. **B**. Distribution of proteins identified from HTEs. **C**. Overlap between proteins identified from LTEs and HTEs. **D**. Overlap between collected proteins identified at the transcript and protein levels. **E**. Number of proteins identified by different LTEs. IDA, IEP, and IMP are the experimental evidence codes in the GO database. **F**. Number of proteins identified by different HTEs. Smaller amounts of data are not shown in the figure.

### Query function and result presentation in iHypoxia

To rapidly access the detailed information on a protein of interest, we recommend that users apply the search function, which is available on both the home and search pages (**Figure 3A**). We also provide examples for each keyword option and allow users to easily experience the search process. Examples include ‘ENSG00000100644’, ‘Q16665’, ‘HIF1A’, and ‘hypoxia inducible factor 1 subunit alpha’, which are specific for the Ensembl ID, UniProt ID, gene name, and protein name, respectively (**Figure 3A**). The database uses a fuzzy query function, and all matched results are listed and displayed in a table at the bottom of the search page, whereas the matched content is highlighted in red (**Figure 3B**). The table showing the results contains basic information, i.e., Ensembl ID, UniProt ID, gene name, protein name and organism, and detailed information for the searched protein can be viewed in the result page by clicking the ‘More’ link (**Figure 3B**).

**Figure 3.**
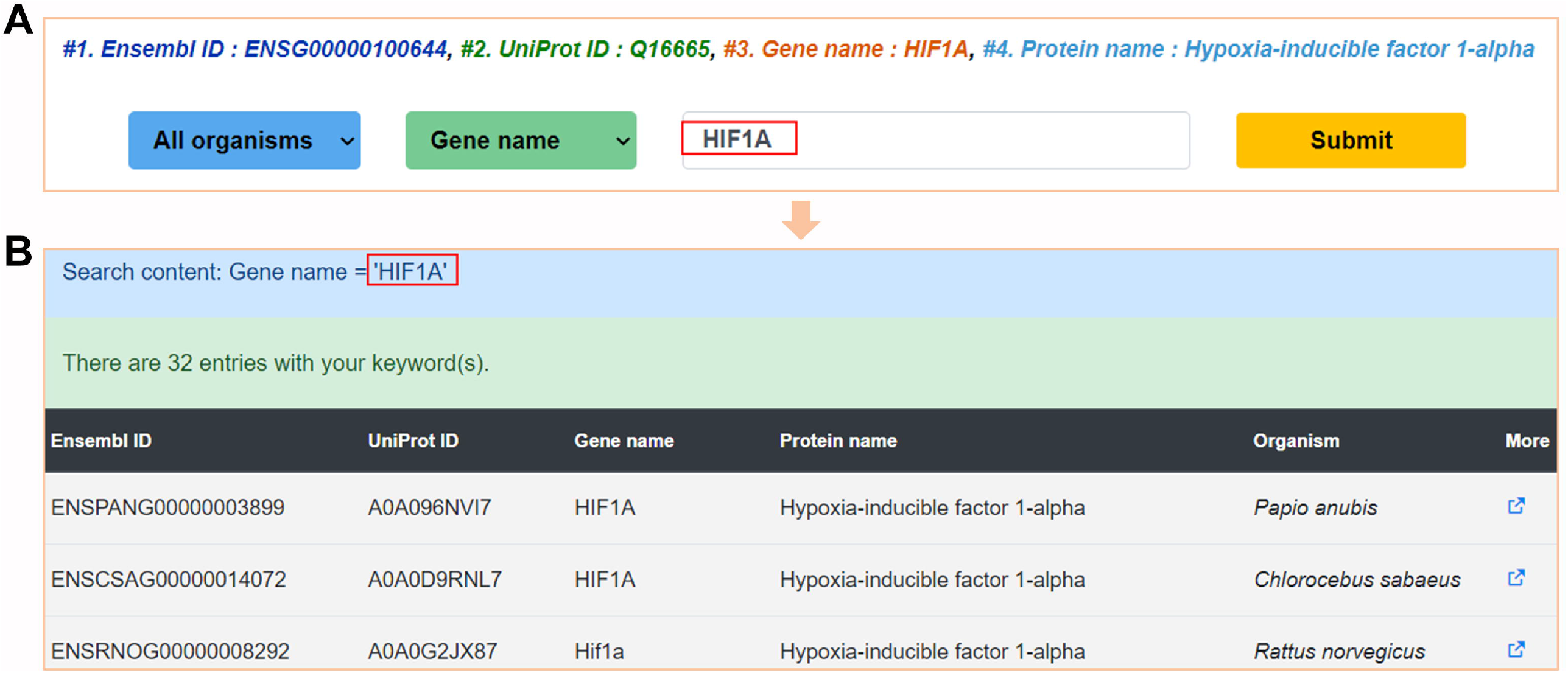
Search option in iHypoxia. **A**. Search options on the home page and search page. **B**. Results returned from a search for a protein.

Because the iHypoxia data were integrated from LTEs, HTEs and an orthologous search, the browser sections are displayed according to these three categories. We first counted the number of proteins in each species identified in LTEs and displayed the result in ‘Statistics for LTE proteins’ (**Figure 4A**). To view the proteins in each organism, users can click the corresponding bar, and a table with the basic information of the Ensembl ID and gene name will be shown on the right (**Figure 4B**). Additionally, users can click the ‘More’ link to view detailed information. For proteins identified in HTEs, because a series of proteins was identified in a single experiment, we preferred to first obtain statistics on the experimental conditions for every organism. The results are displayed in ‘Statistics for HTE conditions’, and users can click one bar to view all the experiments that have been performed with the selected organism (**Figure 4C**). The experimental conditions are shown in a table on the right, which also contains the protein numbers for each condition (**Figure 4D**). Moreover, all proteins obtained under certain conditions in certain organisms are shown at the bottom when users click the ‘View’ button. The table of proteins identified in HTEs includes the Ensembl ID, gene name, log2 ratio, *P*-value and sample type, and detailed information can be accessed through the ‘More’ link (**Figure 4E**). Furthermore, to conveniently browse orthologous proteins, a “Data browse for all organisms” strategy was implemented. In this part, an evolutionary tree showing the phylogenetic relationships of these 50 animals is diagrammed on the right side, and the animals are listed on the left side (**Figure 4F**). Users can click the animal of interest to view orthologs of experimentally identified proteins (**Figure 4G**).

**Figure 4.**
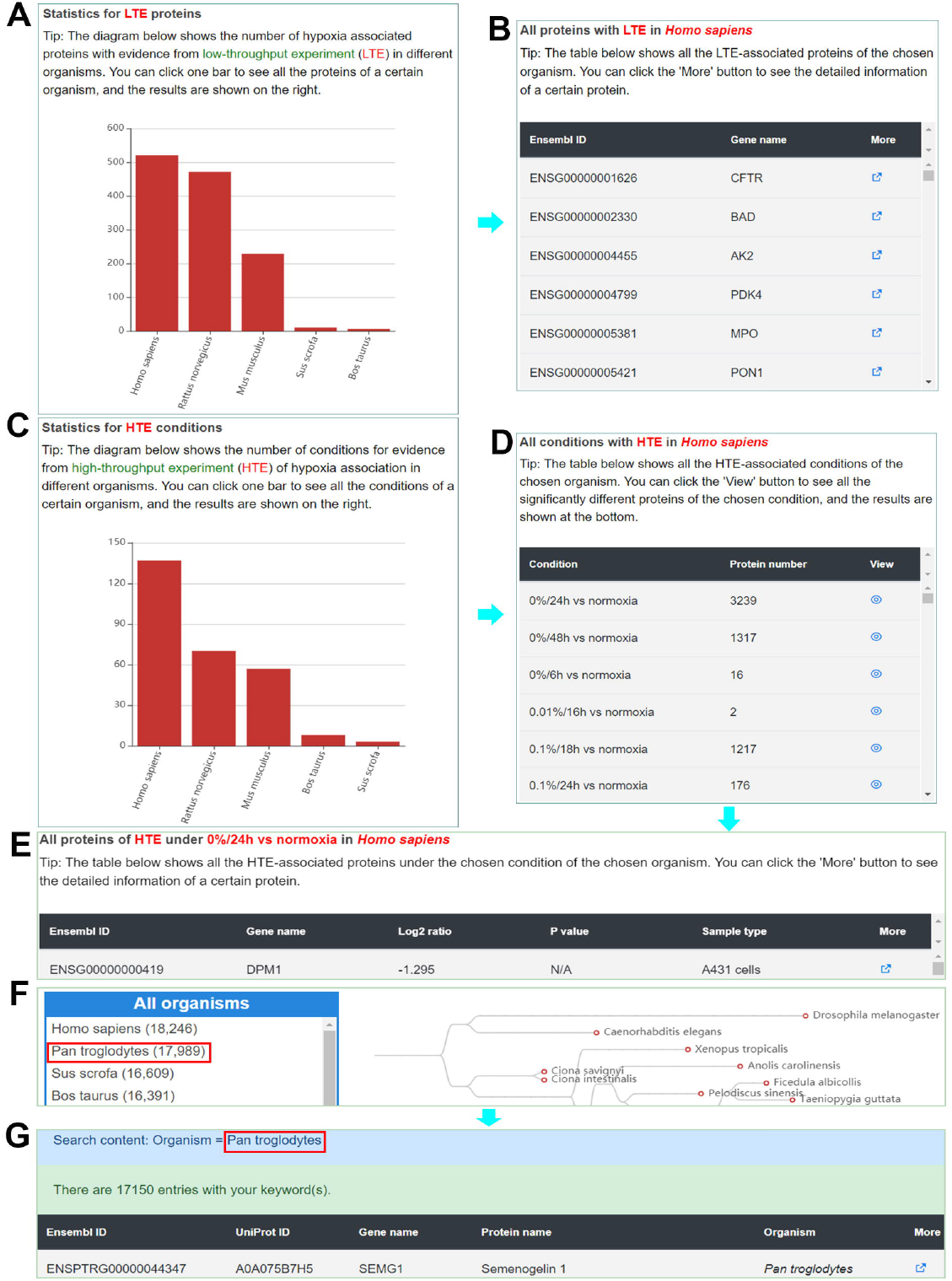
Browsing options in iHypoxia. **A**. Browsing for collected proteins with evidence from LTEs with different organisms. **B**. Search results in a table for proteins identified from LTEs with the chosen organism. **C**. Browsing for proteins with evidence from HTEs with different organisms. **D**. Results in a table obtained with the chosen organism under hypoxic conditions. **E**. Results in a table obtained for proteins identified from HTEs with the chosen organism under the chosen condition. **F**. Browse results by species. **G**. Results in a table for all proteins of the chosen organism.

The result page for a protein consists of nine parts. The first part shows the basic related information of the protein, including the organism, Ensembl ID, UniProt ID, gene name, protein name, protein function and sequence (**Figure 5A**). The other four parts are the expression dynamics of the proteins in response to hypoxia identified in LTEs and HTEs and those obtained from an orthologous search of proteins identified in LTEs and HTEs (**Figures 5B to 5E**). The table of evidence from LTEs contains the experimental conditions, regulation under hypoxic conditions, brief descriptions of the experiments, sample types, evidence level, and detection methods (**Figure 5B**), and the table of HTEs includes the experimental conditions, regulation under hypoxic conditions, log2 ratios, *P*-values, FDRs, sample types, evidence levels, and detection methods (**Figure 5C**). The tables for proteins obtained from an orthologous search showing the evidence from LTEs and HTEs are similar to the tables of evidence from LTEs and HTEs, respectively (**Figures 5D and 5E**). The remaining four parts are ‘Post-translational modification(s)’, ‘Protein-protein interaction(s)’, ‘Subcellular location(s)’ and ‘Drug-target relation(s)’, and these parts correspond to all PTMs we have collected for the searched protein, a network in which the protein interacts with other LTE proteins, subcellular location information and potential inhibitors, respectively (**Figures 5F to 5I**).

**Figure 5.**
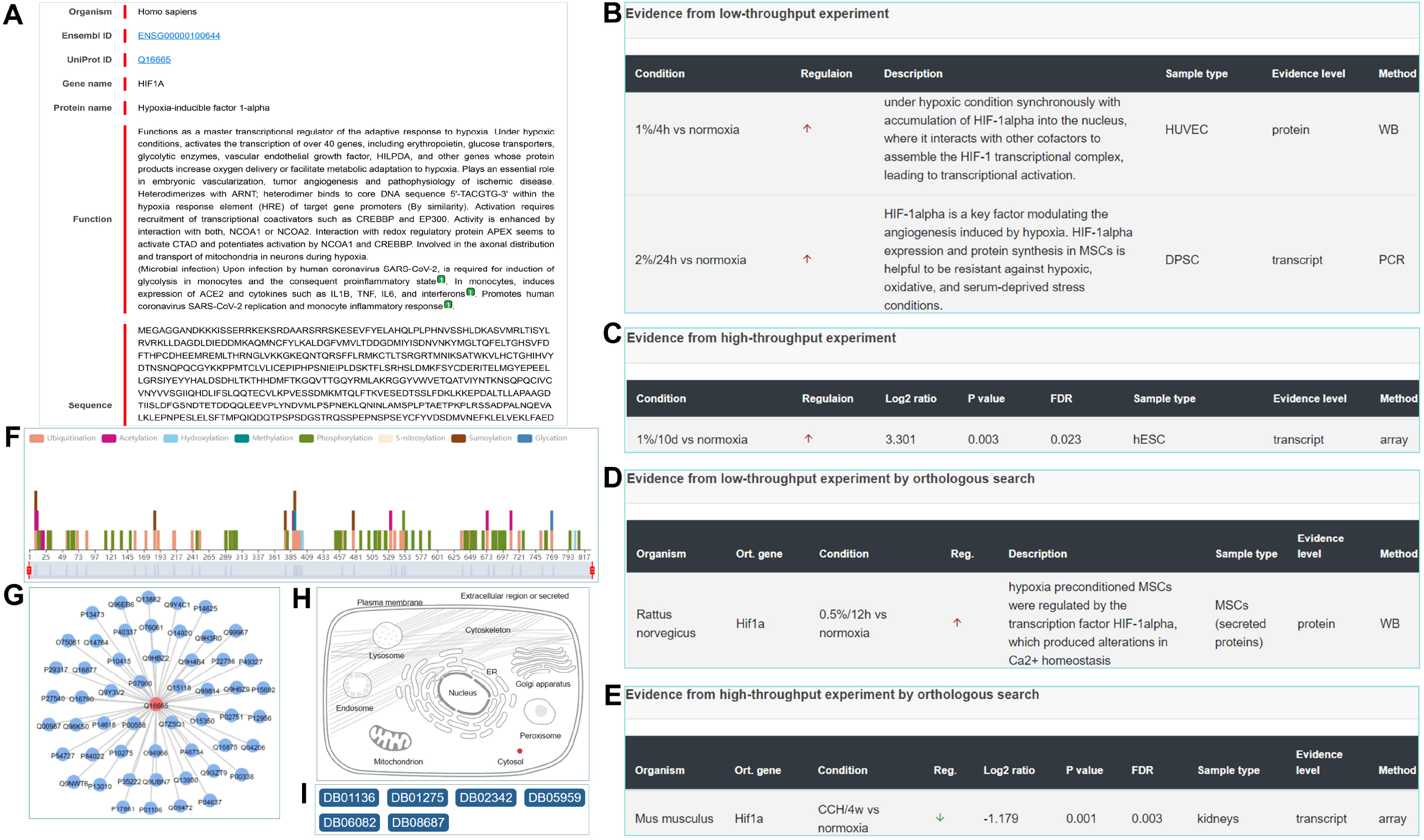
Results page obtained for a searched protein. **A**. Fundamental annotations of a searched protein. **B**. LTE evidence for a searched protein. **C**. HTE evidence for a searched protein. **D**. LTE evidence for an orthologous search for a searched protein. **E**. HTE evidence for an orthologous search for a searched protein. **F**. PTMs for a searched protein. **G**. Protein-protein interactions of a searched protein. **H**. Subcellular location of a searched protein. **I**. Drug-target relationship of a searched protein.

### Functional analyses of collected proteins

To further understand the biological and functional properties of LTE proteins that are differentially expressed in hypoxia, we performed GO [20] and Kyoto Encyclopedia of Genes and Genomes (KEGG) [21] pathway enrichment analyses. For simplicity, only the top 10 significant GO terms of biological processes (*P*-value < 1.73E-32), cellular components (*P*-value < 1.78E-13) and molecular functions (*P*-value < 6.19E-10) and KEGG terms (*P*-value < 3.94E-12) are shown in **Figure 6**. The biological processes were enriched in the regulation of the apoptotic signaling pathway (GO:2001233), response to oxidative stress (GO:0006979), regulation of vasculature development (GO:1901342) and GO terms related to oxygen levels such as response to decreased oxygen levels (GO:0036293), response to hypoxia (GO:0001666), and cellular response to hypoxia (GO:0071456). (**Figure 6**). Many studies have found that the levels of reactive oxygen species (ROS) increase during hypoxia [22], and hypoxia gives rise to apoptotic responses or apoptosis resistance in tumors [23-25]. Systemically, organisms have evolved many vital mechanisms to adapt to hypoxia, and these mechanisms include blood vessel growth, which delivers as much oxygen and nutrients as possible to meet the metabolic demand [4, 26]. With respect to cellular components, most proteins originate from the platelet alpha-granule lumen (GO:0031093), collagen-containing extracellular matrix (GO:0062023), and membrane raft (GO:0045121). The representative molecular functions were 2-oxoglutarate-dependent dioxygenase activity (GO:0016706), monosaccharide binding (GO:0048029), protease binding (GO:0002020) and others. The majority of ‘binding’ terms suggest that proteins that respond to hypoxia play extremely important roles in complex protein interactions. In addition, ‘2-oxoglutarate-dependent dioxygenase activity’ was identified as the most enriched GO molecular function, and dioxygenase can mediate the interaction between HIF-1α and VHL by hydroxylating HIF-1α [8, 9, 27].

**Figure 6.**
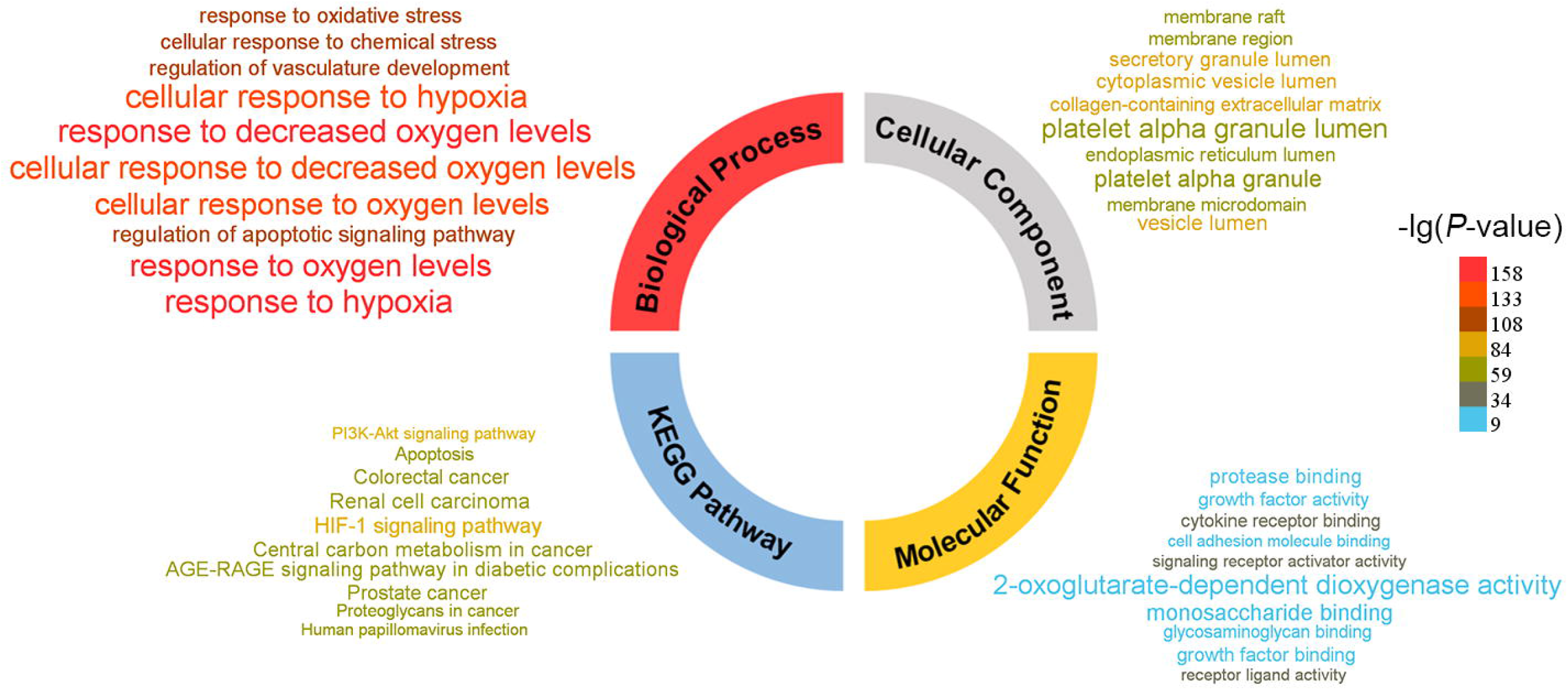
Enrichment of GO and KEGG terms for human LTE proteins. The results were visualized with WocEA (Word cloud for the Enrichment Analysis, version 1.0). The font size denotes the enrichment ratio of each term, and the color denotes the approximate *P*-value.

Hypoxia is a complex condition and a feature of physiological and pathological states. Organisms sense and adapt to hypoxic stress via various and intricate pathways [1]. In addition to the classical and well-known HIF-1 signaling pathway (hsa04066), the top enriched KEGG pathways include renal cell carcinoma (hsa05211) [28], AGE-RAGE signaling pathway in diabetic complications (hsa04933) [29], central carbon metabolism in cancer (hsa05230) [30], colorectal cancer (hsa05210) [31], prostate cancer (hsa05215) [32], apoptosis (hsa04210) [25], proteoglycans in cancer (hsa05205) [33], and the PI3K-Akt signaling pathway (hsa04151) [34]. In particular, many cancer-related pathways are enriched in human proteins identified from LTEs.

We then performed an enrichment analysis with a hypergeometric test by mapping the human proteins identified from LTEs to the druggable proteins acquired from the DrugBank database [35] and oncogenes acquired from the Cancer Gene Census in the Catalog Of Somatic Mutations in Cancer (COSMIC) [36]. As a result, 228 and 54 proteins identified in LTEs were found to be significantly overrepresented in drug targets and cancer genes, respectively (**Table 1**). Therefore, the dysregulation of proteins identified in LTEs may be an essential process in tumorigenesis and development and can be a potential therapeutic target.

PTMs play vitally important roles in regulating a wide range of biological processes. In our KEGG analysis results, one of the remarkably enriched KEGG pathways, the PI3K-Akt signaling pathway, was mainly regulated by phosphorylation. In the iHypoxia database, we provide abundant annotations by taking advantage of the knowledge from several PTM resources (**Figure 1**). We then performed a PTM analysis by mapping known PTM sites from six databases to proteins identified in LTEs. In total, we obtained 35,958 PTM sites on 1,097 substrates, which accounted for 90.29% (1097/1215) of the proteins identified in LTEs (**Figure 7A**). More than 30 modifications were involved, and these included 24,018 phosphorylation sites of 1,056 proteins, 7,124 ubiquitination sites of 637 proteins and 4,662 acetylation sites of 619 proteins (**Figure 7B** and Supplementary Table S2). Different PTMs can engage in crosstalk with each other to synergistically orchestrate specific biological processes, and we found that a fairly large number of proteins identified in LTEs can be regulated by multiple PTMs (**Figure 7C**). Numerous studies have shown that PTMs can be regulated by hypoxia and can also regulate the relevant pathways to respond to hypoxia [37-39].

**Figure 7.**
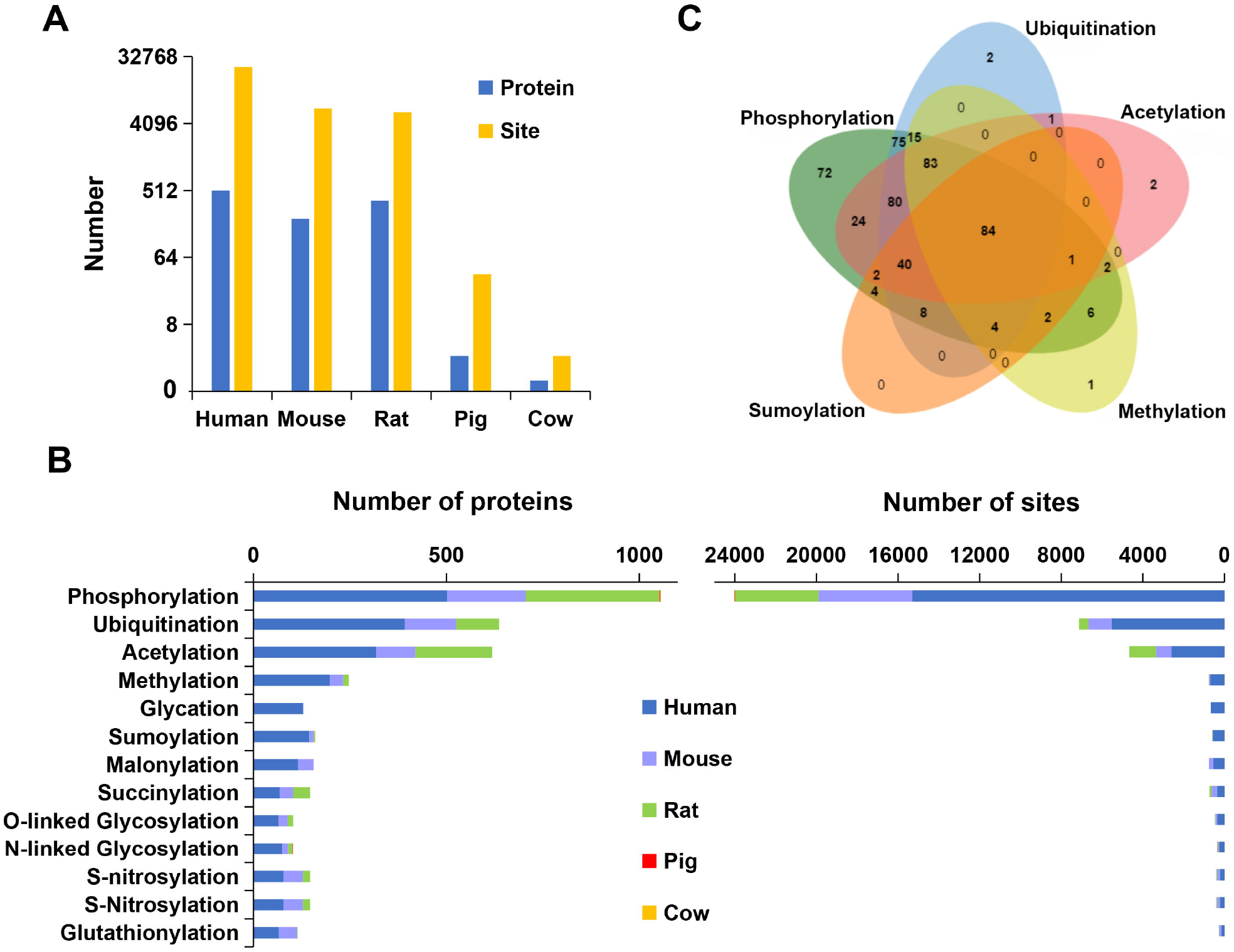
PTM analyses of proteins identified from LTEs. **A**. Distribution of modified proteins and sites. **B**. Distribution of the mapped proteins and sites in terms of phosphorylation, ubiquitination, acetylation and others (sites with fewer than 100 sites are not shown). **C**. Overlap of five major types of PTMs of proteins identified in LTEs.

## Discussion

Oxygen is vital to life on earth. However, hypoxia is a common stress frequently encountered by aerobic species, such as at high altitudes and in aquatic habitats, and underground burrows, and cells/tissues within the body are also exposed to a hypoxic microenvironment. Therefore, mammals have evolved adaptive mechanisms to cope with hypoxia [2-5]. Hypoxia is also associated with human diseases such as cancers, cardiovascular diseases and anemia [15, 16, 40]. The development of high-throughput technologies has greatly enhanced the large-scale identification of proteins whose expression at the transcript or protein levels is modified during hypoxia, and an increasing number of proteins involved in regulating hypoxia have been verified. Considering that the data from these studies are scattered throughout the literature and are difficult to obtain and utilize, a resource for the expression dynamics of proteins in response to hypoxia is needed.

In this study, we developed iHypoxia, an integrated database for the expression dynamics of proteins that respond to hypoxia in animals. At present, iHypoxia contains 1,629 expression events of 1,215 proteins identified in LTEs, 154,953 quantitative expression events of 36,194 proteins identified in HTEs, and 581,763 orthologous proteins. Compared with previous databases, iHypoxia not only contains more data but also has several advanced features. First, iHypoxia is the first integrated animal database focused on the expression dynamics of proteins in response to hypoxia and provides evidence from high- and low-throughput experiments and the transcript and protein levels of five mammals. Moreover, this resource is well annotated and can provide abundant information on proteins, including the hypoxic experimental conditions, expression patterns, samples and other information from the original studies, as well as PTMs, protein-protein interactions, drug-target relations and other information from public databases. Furthermore, the iHypoxia website is convenient and user-friendly, and users can rapidly and accurately find the information of interest or download datasets.

In the future, iHypoxia will be regularly maintained and updated when new hypoxia data are published. In addition, hypoxia not only transcriptionally and translationally activates genes to modulate cellular adaptations but also influences many other processes, such as protein PTMs and the expression of noncoding RNAs [34, 41, 42]. Therefore, data at other expression levels and more species will be integrated to provide a more comprehensive and useful resource to facilitate related research.

## Materials and methods

To develop a comprehensive resource for the expression dynamics of proteins in response to hypoxia in animals, we collected and integrated published (through January 2020) datasets from various sources, such as PubMed, Gene Expression Omnibus (GEO), HypoxiaDB [18], and GO [20]. A workflow of the construction of iHypoxia is depicted in **Figure 1**.

### Curation of protein expression dynamics in response to hypoxia identified in low-throughput experiments and other databases

To obtain the highest quality and validated proteins and their expression patterns in response to hypoxia, we searched for hypoxia-related scientific literature in PubMed using the keyword ‘hypoxia’, and 572 studies ultimately passed our screening process. By carefully reading the literature, we manually extracted the hypoxic expression of proteins identified from LTEs, such as Western blot (WB), Northern blot (NB), PCR, and other experiments. Considering that proteins show different expression patterns in various tissues/cells, in the presence of different oxygen concentrations and in response to different durations of hypoxia, we retrieved as many details as possible, including the hypoxic experimental conditions, expression patterns (upregulation or downregulation), samples (tissues or cells), evidence levels (proteins or transcripts), experimental methods (e.g., WB and NB) and a brief description of the proteins from the original articles. All the protein expression experiments were checked by two data curators. In addition, we integrated data from HypoxiaDB [18] and GO resources [20]. From GO resources, we collected only the proteins whose GO terms were related to hypoxia, as marked by experimental evidence codes, which indicated that the annotation of the gene was directly supported by evidence from an experiment. We then traced the original literature to obtain the expression pattern of the protein, the hypoxic conditions and other information. For the data in HypoxiaDB, detailed information was adopted without further changes.

### Collection of quantitative expression events of proteins in response to hypoxia in high-throughput experiments

Proteomics is a highly effective approach for identifying alterations in protein levels. To identify the expression at the protein level of genes that respond to hypoxia, we searched PubMed with various keywords such as ‘hypoxia AND (proteome OR proteomic)’ and ‘anoxia AND (proteome OR proteomic)’. To control the quality of the data, we manually examined all the literature retrieved and the corresponding supplemental materials. Information pertaining to the hypoxic condition, regulation pattern, *P*-value, sample type and other details was then extracted.

Transcriptomic approaches have been widely used to research transcriptional expression during the process of hypoxic responses, and GEO hosts many expression samples related to processes of hypoxic responses. Therefore, we searched the GEO database with the keywords ‘hypoxia’ and ‘anoxia’. Hypoxia-related microarray expression datasets were then downloaded and analyzed using the limma package [43]. If the fold change of mRNA expression in hypoxic cells/tissues compared with control (normoxia) cells/tissues was greater than 2 and had an adjusted *P-*value <0.05, the protein was included in the iHypoxia database. Detailed information, including the hypoxic experimental conditions, samples, and other details, was also retrieved from the original studies.

After completing the above data collection, the “Retrieve/ID mapping” tool in the UniProt database was used to convert the different identifiers of the collected proteins to obtain a unified Ensembl ID and UniProt ID [44].

### Orthologous detection

To identify the proteins in other animals that may respond to hypoxia, the “Pairwise orthologs” data were downloaded from the OMA database (https://omabrowser.org/oma/current/) [45]. Through screening, we obtained 50 animal ortholog datasets, and the Ensembl ID and UniProt ID were used as identifiers. The proteins were then mapped to the OMA dataset to obtain their orthologs.

### Data obtained from annotation with public databases

To provide abundant annotations for collected proteins, the knowledge obtained from several public databases, including essential information, PTMs, subcellular locations, protein-protein interactions and drug-target information, was integrated.

First, essential information on the proteins, such as the gene name, protein name and sequence, was downloaded from UniProt [46]. PTMs play vital roles in regulating the structure and function of proteins; thus, PTM data from BioGRID [47], dbPTM [48], EPSD [49], PhosphoBase [50], PhosphoSitePlus [51], and PLMD [52] were mapped to the collected proteins. Proteins function in specific subcellular locations, and these locations provide a particular environment and a set of interacting chaperone proteins that are necessary for protein function. Knowledge of the subcellular localization of proteins is important for understanding cellular processes. Therefore, we acquired the subcellular locations from UniProt [46] and protein-protein interactions from BioGRID [47], BioPlex [53], IID [54], IntAct [55], and HuRI [56]. Based on expression studies, some proteins have also been proposed as potential diagnostic or therapeutic biomarkers for human hypoxic diseases. To determine whether the proteins we collected can be used as drug targets, we downloaded the data from DrugBank and matched them [35].

### Implementation of the webserver

All the data in iHypoxia were stored in a MySQL database, and the website was deployed with an Apache server. In the front end, BootStrap, an open-source user interface framework based on HTML, CSS and JavaScript, was used to build the basic layout of the website. JavaScript and its tool library JQuery were used to implement the interactive functions. The visualization of the data statistics and PTMs was achieved with Echarts, whereas the network of protein-protein interactions was displayed using Cytoscape.js. Data processing in the back end was performed through PHP, and the interaction between the front and back ends was achieved through AJAX. Moreover, to provide a stable and adaptive service, we tested the iHypoxia website using various web browsers, including Mozilla Firefox, Google Chrome, and Internet Explorer.

## Data availability

The data in this study is available at http://ihypoxia.omicsbio.info.

## CRediT authorship contribution statement

**Ze-Xian Liu**: Conceptualization, Methodology, Investigation, Writing - review & editing, Project administration, Funding acquisition. **Panqin Wang**: Data Curation, Investigation, Writing - original draft. **Qingfeng Zhang**: Data Curation, Software, Writing - original draft. **Shihua Li**: Data Curation, Validation. **Yuxin Zhang**: Data Curation, Validation. **Yutong Guo**: Data Curation, Validation. **Chongchong Jia**: Data Curation, Validation. **Tian Shao**: Data Curation, Validation. **Lin Li**: Validation. **Han Cheng**: Conceptualization, Writing - review & editing, Funding acquisition. **Zhenlong Wang**: Conceptualization, Writing - review & editing, Funding acquisition.

## Acknowledgements

This work was supported by grants from the Natural Science Foundation of China [U2004152 to ZL.W., 81972239 and 91953123 to ZX.L., 31601067 to H.C.], Program for Guangdong Introducing Innovative and Entrepreneurial Teams [2017ZT07S096 to ZX.L.], and Tip-Top Scientific and Technical Innovative Youth Talents of Guangdong special support program [2019TQ05Y351 to ZX.L.].

## Tables

**Table 1** Enrichment analyses of drug targets and cancer genes for human LTE proteins.

## Supplementary material

**Supplementary Table S1** Numbers of experimentally or computationally identified proteins in 50 animals in the iHypoxia database.

**Supplementary Table S2** Distribution of substrates and different PTM sites of collected proteins identified from LTEs.

