## Supplementary material for "iHypoxia: an integrative database of the expression dynamics of proteins in response to hypoxia in animals": Table 1

**Table 1 Enrichment analyses of drug targets and cancer genes for human LTE proteins**

|  | **Hypoxia** | | **Proteome** | | **E-ratio** | ***P*-value** |
| --- | --- | --- | --- | --- | --- | --- |
|  | **m** | **n** | **M** | **N** |  |  |
| Drug target | 228 | 508 | 2895 | 22784 | 3.53 | 1.37E-73 |
| Cancer gene | 54 | 508 | 711 | 22784 | 3.41 | 2.89E-15 |
